# Single-cell Genome-and-Transcriptome sequencing without upfront whole-genome amplification reveals cell state plasticity of melanoma subclones

**DOI:** 10.1101/2023.01.13.521174

**Authors:** Koen Theunis, Sebastiaan Vanuytven, Florian Rambow, Daniel Brown, Michiel Van Der Haegen, Oskar Marin-Bejar, Aljosja Rogiers, Nina Van Raemdonck, Eleonora Leucci, Jonas Demeulemeester, Alejandro Sifrim, Jean-Christophe Marine, Thierry Voet

## Abstract

Single-cell multi-omics methods are enabling the study of cell state diversity, which is largely determined by the interplay of the genome, epigenome, and transcriptome. Here, we describe Gtag&T-seq, a genome-and-transcriptome sequencing (G&T-seq) protocol of the same single cells that omits whole-genome amplification (WGA) by using direct genomic tagmentation (Gtag). Gtag drastically decreases the cost and improves coverage uniformity at both the single-cell and pseudo-bulk level when compared to WGA-based G&T-seq. We also show that transcriptome-based DNA copy number inference has limited resolution and accuracy, underlining the importance of affordable multi-omic approaches. Moreover, applying Gtag&T-seq to a melanoma xenograft model before treatment and at minimal residual disease revealed differential cell state plasticity and treatment response between cancer subclones. In summary, Gtag&T-seq is a low-cost and accurate single-cell multi-omics method enabling the exploration of genetic alterations and their functional consequences in single cells at scale.

## INTRODUCTION

Single-cell genomics is key to study genetic heterogeneity in cells of developing and ageing organisms in health and disease. Compared to conventional bulk DNA sequencing, these methods offer enhanced resolution to characterise the mutational processes operative in tissues and organs (Bian et al., 2018; Navin and Hicks, 2011; Pellegrino et al., 2018). Leveraging the detected somatic mutations, the phylogeny of the cells and tissues can be reconstructed (Bae et al., 2018; Eirew et al., 2015; Gao et al., 2016; Kim et al., 2018; Navin et al., 2011; Wang et al., 2014), increasing our understanding of the developmental and evolutionary mechanisms of (diseased) tissues as well as of the aetiological role of the acquired mutations in the phenotype (Cai et al., 2014; Lodato et al., 2018).

The majority of single-cell genome sequencing methods require whole-genome amplification (WGA) to preamplify the genomic DNA (gDNA) and obtain enough material for preparing a sequencing library. In general, WGA methods apply either Polymerase Chain Reaction (PCR, *e.g.* DOP-PCR, LM-PCR, Ampli1) (Ferrarini et al., 2018; Klein et al., 1999; Telenius et al., 1992), multiple displacement amplification (MDA; *e.g.* RepliG, GenomiPhi) (Dean et al., 2002), or a combination of both (*e.g.* MALBAC, picoPlex) (Langmore, 2002; Zong et al., 2012). Amplification, however, introduces coverage bias and nucleotide errors due to buffer and polymerase artefacts. Additionally, the process of preamplification followed by library preparation is low throughput, labour intensive and costly. Recently, tagmentation-based methods were reported, preparing sequencing libraries from single-cell gDNA without upfront WGA (Laks et al., 2019; Rohrback et al., 2018; Vitak et al., 2017; Xi et al., 2017; Zachariadis et al., 2020), and providing a more uniform representation of the cell’s genome, while enabling a higher throughput.

Single-cell multi-omics technologies have been developed to simultaneously assess the genome, epigenome, transcriptome, and/or selected proteins of single cells by either joint or parallel processing of the molecular hierarchy, or by data analysis methodologies (Angermueller et al., 2016; Cao et al., 2018; Clark et al., 2018; Dey et al., 2015; Hou et al., 2016; Li et al., 2015; Macaulay et al., 2015; Pott, 2017; Zachariadis et al., 2020). Methods for the joint processing of gDNA and RNA of single cells either use a single-tube gDNA-and-mRNA (DR-seq) preamplification followed by separate library preparation (Dey et al., 2015) or rely on their physical separation before their parallel amplification (Macaulay et al., 2015; Han et al., 2014; van Strijp et al., 2017; Zachariadis et al., 2020). DR-seq preamplifies gDNA and mRNA simultaneously, minimizing nucleic acid loss, but complicating data analysis and interpretation due to the presence of RNA-derived reads in the gDNA-seq data. Physical separation of mRNA and gDNA, as in Genome-and-Transcriptome sequencing (G&T-seq) (Macaulay et al., 2015) and DNTR-seq (Zachariadis et al., 2020) yields a more flexible protocol and allows the RNA- and gDNA-seq protocols to be adapted to the researcher’s needs. Up to now, gDNA sequencing in G&T-seq necessitates WGA techniques, making large-scale G&T-seq cost prohibitive and time consuming. Other groups have studied the interplay between genome and transcriptome, by either inferring copy number alterations (CNA) from gene expression profiles (Patel et al., 2014; Tirosh et al., 2016) or integrating separate single-cell gDNA and RNA datasets (Campbell et al., 2019; McCarthy et al., 2020). While these approaches have led to new insights into tumour heterogeneity, tumour progression, as well as therapeutic strategies, only direct multi-omics approaches allow genotype-phenotype relations to be unambiguously ascertained (Bock et al., 2016; Chappell et al., 2018; Macaulay et al., 2017).

Here, we present Gtag&T-sequencing, a method that enables genome-and-transcriptome sequencing of single cells without upfront WGA, enhancing throughput while minimising coverage bias, amplification noise, and cost. Gtag&T-seq of a patient-derived xenograft (PDX) melanoma model highlights the advantages of genome-based over transcriptome-based CNA inference, and the transcriptional effects of complex genomic alterations. We construct a DNA-based cell lineage tree annotated with RNA-based cell type and state information from the same cells, providing unique insights in the role of genetic and non-genetic factors as well as their interplay during tumour evolution under therapeutic pressure.

## RESULTS

### WGA-free parallel genome and transcriptome sequencing of single cells

Gtag&T-seq is based on genome-and-transcriptome (G&T) sequencing (**Fig. 1A**), developed by Macaulay *et al*. (Macaulay et al., 2015, 2016). Following physical separation of the DNA and mRNA of the same cell, Gtag&T-seq applies tagmentation to produce a gDNA sequencing library directly from the cell’s genome, instead of preamplifying it with WGA (**Fig. 1A**). Following tagmentation, PCR adds cell-specific barcodes and sequencing adapters to enable multiplexed low coverage sequencing and cost-effective multi-modal analysis of single cells (**Fig. 1B**).

**Figure 1.**
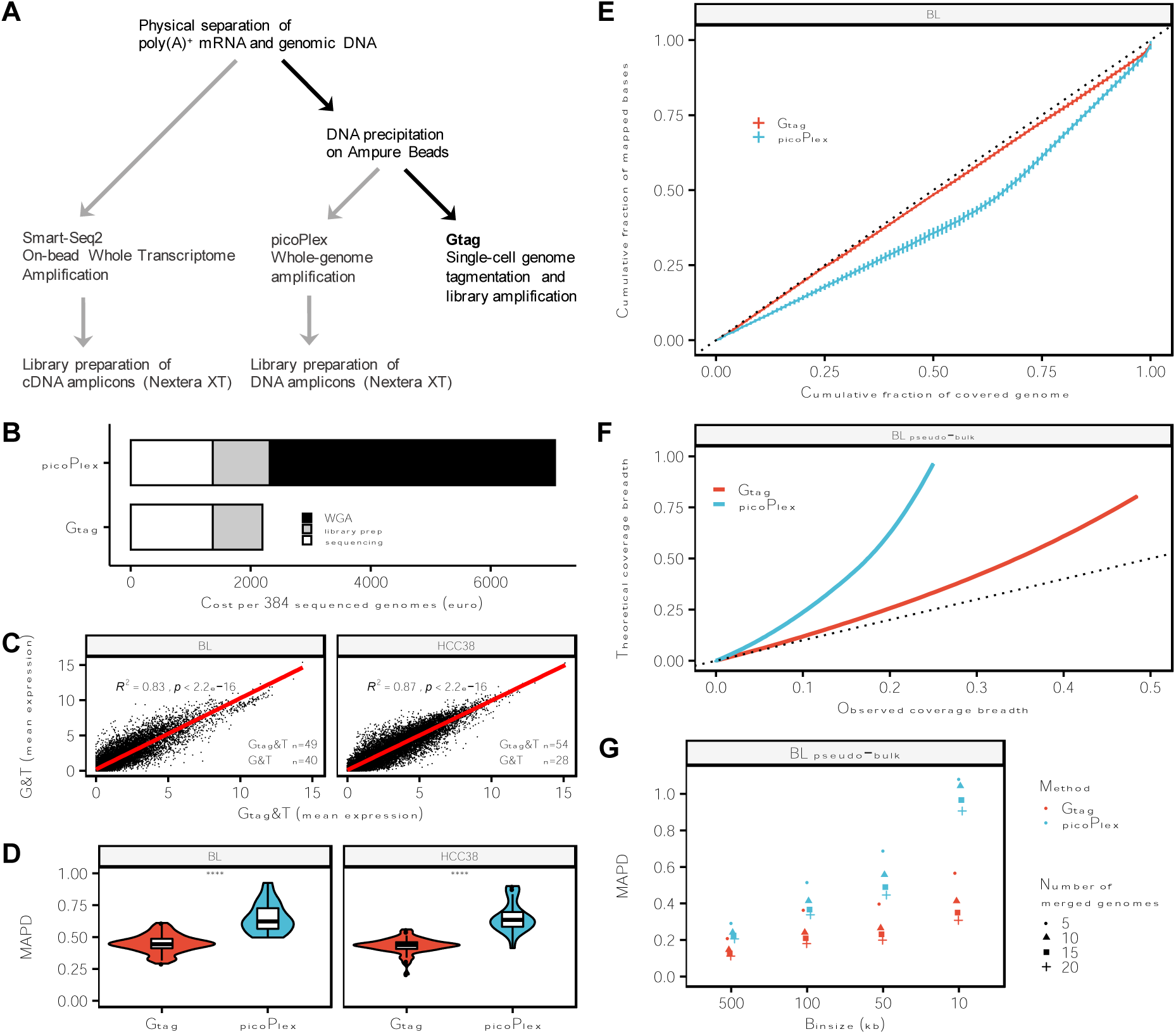
Gtag&T is a cost-effective method for joint analysis of DNA and RNA of single cells. **A** General workflow of genome-and-transcriptome separation followed by Gtag (in black) or conventional G&T-seq with picoPlex and Smart-Seq2 (in grey). **B** Cost of 384 cells processed either by picoPlex (1/2 volumes) and Nextera XT (1/10^th^ volumes) or by Gtag after G&T-separation and sequencing performed on one HiSeq4000 lane. **C** Spearman correlation of the mean expression per gene after conventional G&T-seq *versus* Gtag&T-seq for the HCC38 and matched normal (BL) cell line. **D** Median absolute pairwise difference (MAPD) of logR values between consecutive genomic bins, for HCC38 and BL single-cell DNA samples processed by Gtag or picoPlex, and analysed using genomic bins of 500 kb. **E** Lorenz curves showing the mean coverage uniformity of single-cell genomes. The dotted line represents perfect uniformity. Vertical lines on the Lorenz curves represent the standard deviations from the mean. **F** Observed genome coverage of merged Gtag and picoPlex genomes for a given theoretical genome coverage. The coverage indicated with the dotted line is the coverage maximally attainable assuming no overlapping reads. For each method, increasing amounts of single-cell genomes are merged, plotted, and a local regression is performed. **G** MAPD values for Gtag and picoPlex genomes obtained by merging either 5, 10, 15, or 20 single-cell genomes for different genomic bin sizes. The centreline, top and bottom of the boxplots represent respectively the median, 25th and 75th percentile and whiskers are 1.5 × IQR. ****: p <= 0.0001 (Wilcoxon test).

The performance of Gtag&T-seq was evaluated against conventional G&T-seq (Macaulay et al., 2015), using picoPlex for WGA, for both the HCC38 cancer cell line and its matched normal cell line (BL). PicoPlex was chosen for its proven reproducibility and high accuracy to detect DNA copy numbers (Chen et al., 2018; Deleye et al., 2017; Zhang et al., 2017). At the RNA level, the datasets were comparable, highlighted by the high correlation of the mean expression per gene for both cell lines (BL, *R^2^* = 0.83; HCC38, *R^2^* = 0.87; **Fig. 1C** and **Supplemental Fig. S1A,B**). To allow for a fair comparison of the genome sequences, gDNA reads of each cell were down-sampled to a maximum of 400,000 reads before duplicate removal (Methods and **Supplemental Data S1**), a depth obtainable by sequencing 384 libraries on one Illumina HiSeq4000 lane. Reliable genetic variant detection from single-cell gDNA data is largely dependent on the noise and coverage uniformity attained by the method. Genomic readouts were significantly less noisy for Gtag (BL, *p ≤* 0.0001; HCC38, *p* ≤ 0.0001) as assessed by median absolute pairwise difference (MAPD) (**Fig. 1D**). In addition, Gtag provided improved coverage uniformity, discernible from Lorenz curves (**Fig. 1E** and **Supplemental Fig. S1C,D**) and Gini index comparison (μ_Gtag_ = 0.20 *versus* μ_picoPlex_= 0.27; *p ≤* 0.0001) of individual BL genomes.

Low-depth single-cell genomes can be pooled to derive pseudo-bulk genomes, enabling refinement of genomic variant characterization (Laks et al., 2019; Zahn et al., 2017). To investigate differences in performance resulting from omitting WGA, we compared coverage breadth, uniformity, and noise after merging single-cell genomes (HCC38-BL) *in silico* for both multi-omics methods. Plotting the theoretical *versus* the observed coverage breadth for increasing amounts of pooled single-cell genomes (**Fig. 1F**) showed a rapid saturation of coverage breadth for picoPlex, which is most likely caused by limited random priming during WGA. Pseudo-bulks can be further leveraged to map DNA breakpoints more precisely, conditional on smaller bin sizes not exacerbating noise. We observed that picoPlex suffered from inflated MAPD scores in comparison to Gtag for pseudo-bulks analysed with smaller bin sizes (**Fig. 1G**). In addition, a more even coverage uniformity was obtained for Gtag 20-cell pseudo-bulk genomes as evidenced by Gini indexes (μ_Gtag_ = 0.18, *versus* μ_picoPlex_= 0.42; *p ≤* 0.0001).

### Single-cell and pseudo-bulk analysis of a human melanoma PDX model

To study genetic subclonal dynamics, cellular phenotypic heterogeneity –including phenotype switching or a more continuous plasticity (Shen and Clairambault, 2020)– as well as genotypic-phenotypic interactions during cancer treatment at single-cell resolution, we processed 165 and 176 single cells from a melanoma PDX model using Gtag&T- and G&T-seq, respectively. The MEL006 model was established from a patient with a BRAF^V600E^ mutant melanoma that had an almost complete response to combined dabrafenib (BRAF^V600E^ inhibitor) and trametinib (MEK inhibitor) therapy (Rambow et al., 2018). Single cells from the PDX model were collected before treatment (T0) and at minimal residual disease (MRD; T28), when most of the tumour cells are eradicated by the therapy. However, a small subset of the cancer cells persists in a drug-tolerant state, providing a substrate for relapse. The molecular mechanisms underpinning drug tolerance and/or resistance may comprise genetic as well as non-genetic plasticity factors. A deeper understanding of these genetic and non-genetic mechanisms is required for the design of longer-lasting combination treatments.

After removing low quality genomes (Methods and **Supplemental Data S2**), 150 single-cell Gtag genomes were compared to 173 single-cell picoPlex genomes. DNA copy number profiles were called using genomic bins of 500 kb, revealing a highly rearranged tumour cell population (**Fig. 2A**) with an average ploidy of 3.5, indicative of an early whole-genome doubling event, coherent with previous bulk sequencing observations (Marin-Bejar et al., 2021).

**Figure 2.**
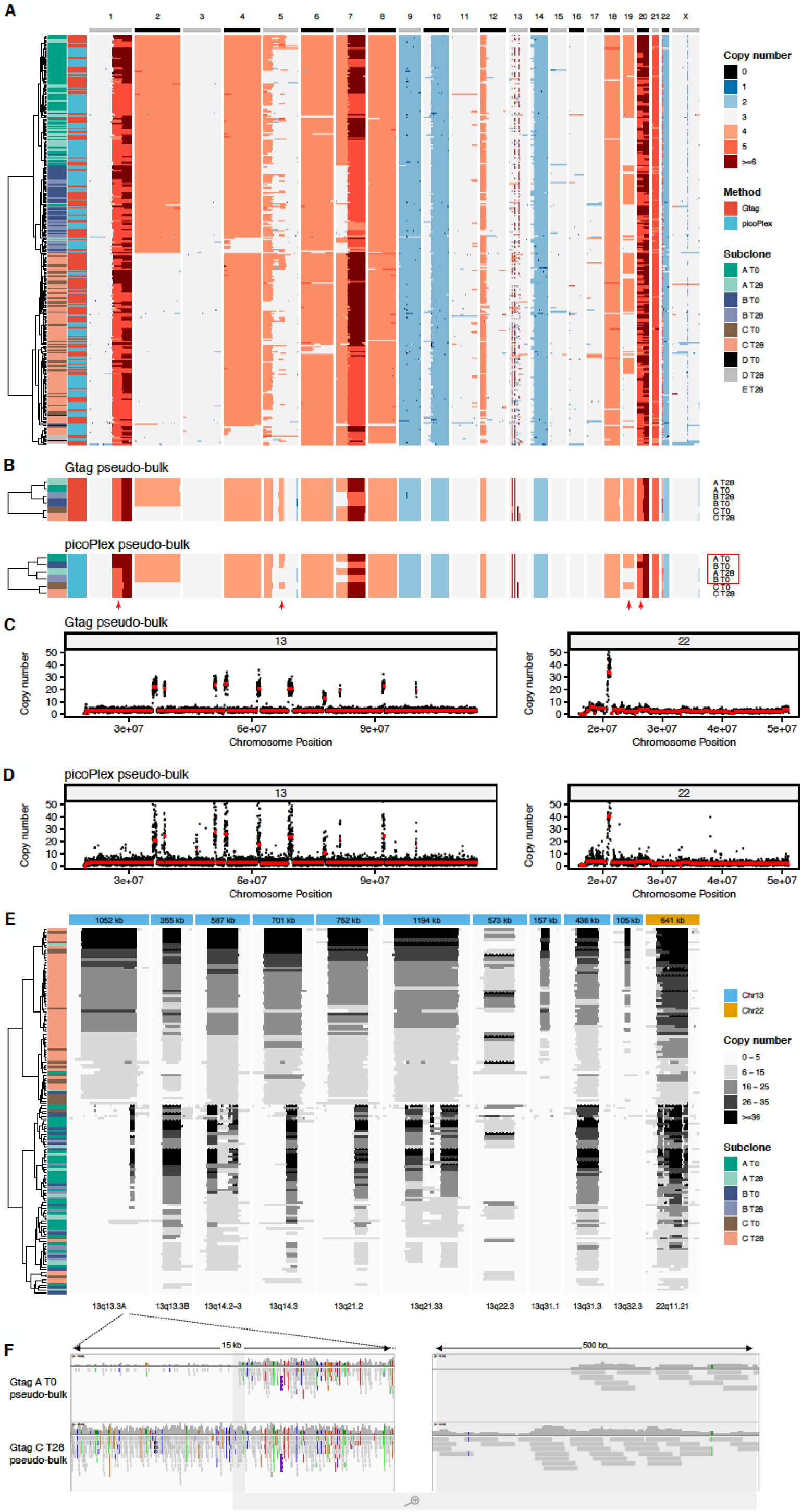
Gtag improves breakpoint and copy number calling in a melanoma xenograft model. **A** Genome-wide copy number heatmap of 323 single cells from a patient-derived xenograft (PDX) model of human melanoma. Columns correspond to chromosomes (divided into genomic bins of 500 kb) and rows to single cells (*n* = 323). All libraries were down-sampled to 400,000 reads before duplicate removal. **B** DNA copy number heatmap of pseudo-bulk genomes (genomes are merged per method, subclone, and time point). Gtag and picoPlex pseudo-bulk genomes are plotted separately. Red arrows indicate incorrect CNA detection resulting in incorrect clustering of subclone A and B (red rectangle) based on pseudo-bulk genomes. In **A** and **B** cells are annotated for their subclone assignment and time point of isolation, as well as the methodology used for DNA processing. **C**, **D** Copy-number plots from respectively Gtag and picoPlex pseudo-bulk genomes (subclone A at T0) of chr13 (left panel) and chr22 (right panel). Black dots represent genomic bins of 10 kb, and the red line represents the segmented integer DNA copy number. **E** Heatmap of the focal amplifications on chr13 and chr22 (annotated in top row in blue and orange respectively). The approximate size of the focal amplification is depicted at the top. DNA copy numbers are indicated in greyscale. **F** View, adapted from IGV, of read distribution in Gtag pseudo-bulk genomes (A T0 *versus* C T28). A breakpoint of subclone A is represented at kilobase resolution (left panel) and base pair resolution (right panel).

Five distinct subclones (A–E; **Fig. 2A**) were identified using hierarchical clustering on Canberra distance. Subclone D (*n* = 8) and E (*n* = 1) contained a small number of cells and were omitted from downstream analyses. The three major subclones differed by the presence of an additional copy of chr2 in A and B, and a loss of one copy of chr7 in subclone B. Within each subclone, single cells displayed similar copy number profiles (mean pairwise R: A= 0.86, B = 0.86, C= 0.78), indicative of stable clonal expansions. To refine the CNA states and breakpoints, pseudo-bulk genomes were created per method for each of the subclones (A-C) at both time points. Similar to our observations with cell lines, WGA resulted in increased noise and decreased coverage uniformity in pseudo-bulk genomes (**Supplemental Fig. S2F**), leading to incorrect clustering of the picoPlex-derived pseudo-bulks (**Fig. 2B**). As a consequence, Gtag pseudo-bulk genomes allowed for a more detailed dissection of focal CNAs (**Fig. 2C,D**).

### Detection of subclone-specific focal amplifications at near base-pair resolution using Gtag

Single-cell and pseudo-bulk CNA profiling with 500 kb bins (**Fig. 2A-D**) indicated the existence of highly amplified loci on chr13 and chr22. The elevated levels of amplification in these regions combined with the lower noise and higher uniformity of Gtag genomes allowed us to further refine the breakpoints using 10 kb bins. In total, 11 focal amplifications ranging in size from 0.12 to 1.20 Mb were found: 10 on chr13 (q13.3A, q13.3B, q14.2, q14.3, q21.2, q21.33, q22.3, q31.1, q31.3, q32.3) and 1 on chr22 (q11.21). GISTIC2 analysis of 367 skin cutaneous melanoma samples recently revealed that 13q and 22q arms are frequently amplified (Broad Institute TCGA Genome Data Analysis Center, 2016). Moreover, a 22q11.21 focal amplification is found in 10% of the CCLE skin cancer cell lines (*e.g.* Hs294-T and COLO679 BRAF(V600) mutant cell lines), 22q11.21 was also identified as recurrently amplified in a pan-cancer analysis (Zack et al., 2013), and was recently associated with inferior survival in acral melanoma (Farshidfar et al., 2022).

Subclonal differences in the presence, size and copy number were observed for the majority of the focal amplifications on chr13 (**Fig. 2E**). Subclone C was characterised by the presence of 13q31.1 and 13q32.3 amplicons, as well as different breakpoints for 4 focal amplifications (*e.g.* 13q13.3A, 13q14.3, 13q21.2, and 13q21.33; **Fig. 2E**). Furthermore, a small number of cells belonging to subclone A and B showed additional breakpoints in 13q14.2-3 and 13q21.33, resulting in multiple smaller amplicons (**Fig. 2E**) as well as small alterations in 22q11.21 that were only detected in Gtag genomes and not in picoPlex genomes, but are confirmed using pseudo-bulk genomes as well as transcriptomic analysis (**Supplemental Fig. S3**). Within subclones, however, most of the breakpoints were conserved (mean pairwise R: A= 0.73, B = 0.72, C= 0.71).

The copy number of the majority of chr13 amplicons was significantly correlated with the amplification level of 22q11.21, suggesting that these amplicons are co-amplified (**Supplemental Fig. S4**). The absence of correlation for amplicon 13q22.3 is explained by its absence in half of the cells irrespective of clonal context. All aforementioned subclonal differences were also detected in pseudo-bulk genomes and remained after downsampling to the same depth (**Fig. 2C-D** and **Supplemental Fig. S5**). Furthermore, these pseudo-bulk genomes allowed us to pinpoint the breakpoint of the focal amplifications to near base-pair resolution (**Fig. 2F**).

Although full reconstruction of the amplicon structure is challenging from single-end sequencing data, we were able to make two key observations. First and foremost, the subclonal organization of these focal amplifications supports the existence of at least 3 major genomic subpopulations in this tumour, with subclones A and B closely related. Secondly, the heterogeneity in the copy number of the amplicons indicates that these are dynamic instead of static DNA entities.

### Transcriptome-based DNA copy number inference has limited accuracy and fails to detect focal amplifications

In order to address the added value of direct genome measurements along with transcriptome profiling we first set out to identify the three major genomic subclones from the single-cell transcriptome data. Of the G(tag)&T single-cell transcriptomes, 256 passed our quality thresholds (Methods and **Supplemental Data S2**) and were used for downstream analysis. We inferred CNAs using averaged gene expression patterns with normal human melanocytes as a reference using inferCNV (Patel et al., 2014). Comparison to the matching gDNA-derived DNA copy number profiles of the same cells, revealed an average sensitivity and specificity of respectively, 48% and 90% (Methods). InferCNV failed to detect the copy number of the whole-chromosome gains for chromosomes 4, 6, 8 and 18 for all cells, as well as chr2 for cells belonging to subclones A and B (**Fig. 3A**). Instead, we obtained the correct copy number for smaller regions of these chromosomes (**Fig. 3A**), suggesting that not all genes are affected to the same degree by genomic imbalances. Furthermore, the focal amplifications on chromosomes 13 and 22 and the p-ter amplification of chr12 were not detected. These shortcomings notwithstanding, we trained a Support Vector Machine (SVM) model on our single-cell genome-and-transcriptome data that achieved a mean classification accuracy of 0.72 [95% CI 0.58-0.84]. Although we were able to identify the three subclones to some degree, we found a high misclassification rate of subclone B to A. Combining subclones A and B, increased the accuracy of the model to 0.92 [95% CI 0.81 - 0.98]. This highlights the need for direct multi-omics to accurately dissect both genomic evolution and transcriptome plasticity in full.

**Figure 3.**
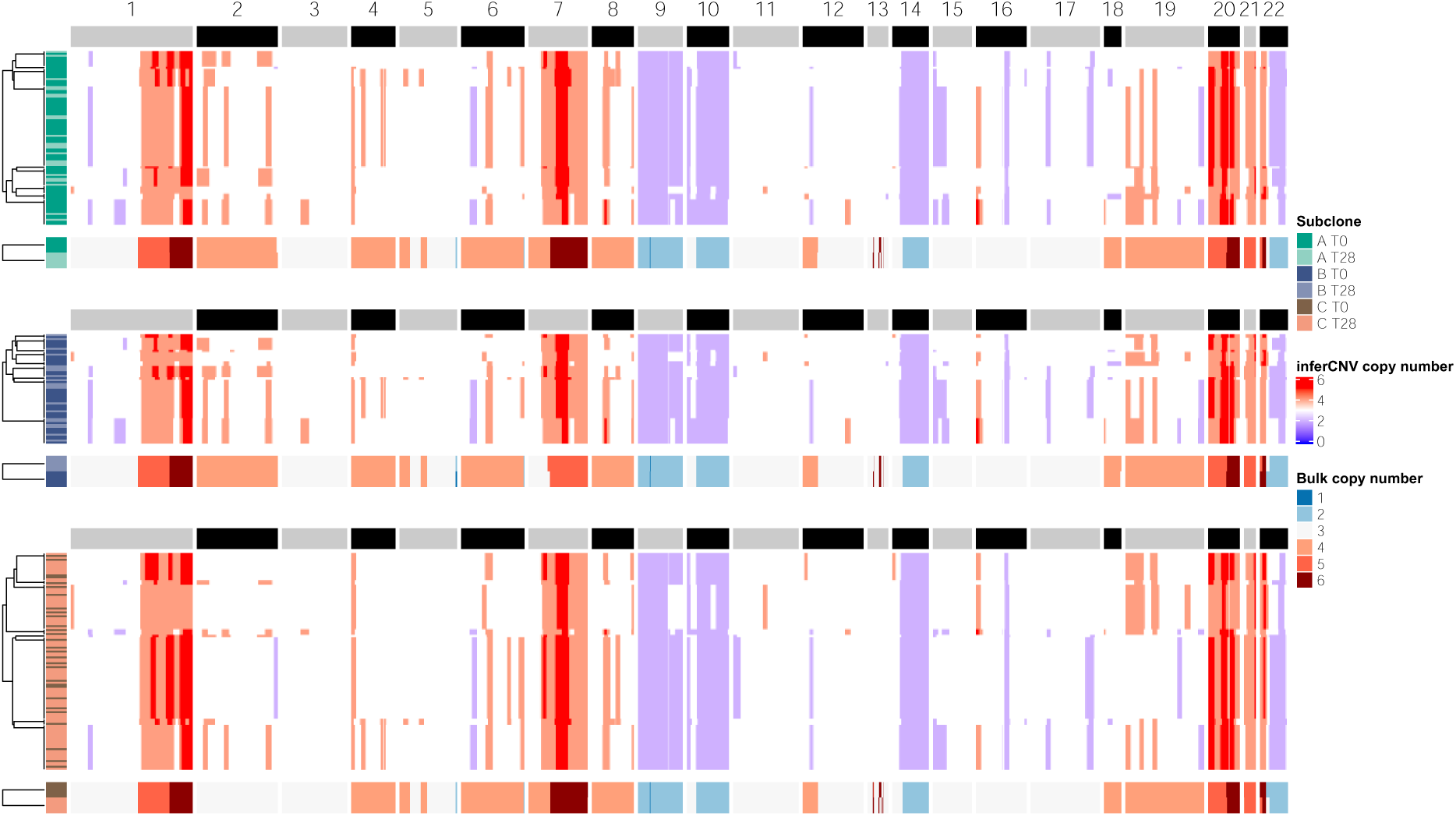
InferCNV is unable to accurately reconstruct CNAs and discern the focal amplifications on chr13 and chr22. Copy number alteration heatmap obtained using inferCNV on the transcriptome data of both the G&T-seq and Gtag&T-seq datasets. Single cells are annotated according to their subclone assignment and time point of isolation. The reference Gtag&T pseudo-bulk copy number profiles per subclone are depicted underneath the inferCNV results for each subclone. All copy numbers were capped at 6 to allow for a fair comparison between the methods.

### Differential expression analysis reveals subtle effects of subclonal chromosomal alterations

Previously, single-cell RNA-seq of the same PDX melanoma model exposed to BRAFi/MEKi treatment, identified four inducible drug-tolerant cell states: a “starved” (starved-like melanoma cell, SMC) state that could either differentiate into a “pigmented” state, or de-differentiate into an “invasive/mesenchymal-like (or undifferentiated)” state or a “neural crest stem cell (NCSC)-like” state (Rambow et al., 2018). To investigate the effects of the genomic alterations on the cell’s transcriptome, and the aforementioned drug-tolerant states, we performed pairwise differential geneexpression analysis between the three genomic subclones at each time point. In total, only 14 differentially expressed genes were identified (FDR < 0.05) (**Fig. 4A**). At both T0 and T28, no difference was found between subclones A and B, indicating that the loss of one copy of chr7 has only subtle effects on the transcriptomes of cells belonging to subclone B. This observation agrees with the difficulties of the SVM model to correctly classify subclone B based on the CNAs obtained with inferCNV. In contrast, we found that loss of chr2 in subclone C caused lower expression of *PCBP1* and *ARPC2* compared to A, and *PCBP1* and *OST4* to subclone B. At T28, *AQP1* (chr7) and *L1CAM* (chrX), both markers of the NCSC state, showed increased expression in subclone A and B respectively, while *CD36* (chr7), a marker for the SMC state, was expressed at higher levels in subclone C compared to subclone A. Three differentially expressed genes were located on the focal amplicons of chr13 and chr22. At both T0 and T28, *THAP7* (22q11.21) was found to be higher expressed in subclone C, while *LZTR1* (22q11.21) was additionally found at T28. Lastly, *GPC5* (13q31.3) had a higher expression in cells belonging to subclone B at T28. These observations suggest the presence of subclonal differences in expression levels of genes located on the focal amplifications, or alternatively, differences in the copy number of the amplicons influencing the gene expression between subclones A, B and C.

**Figure 4.**
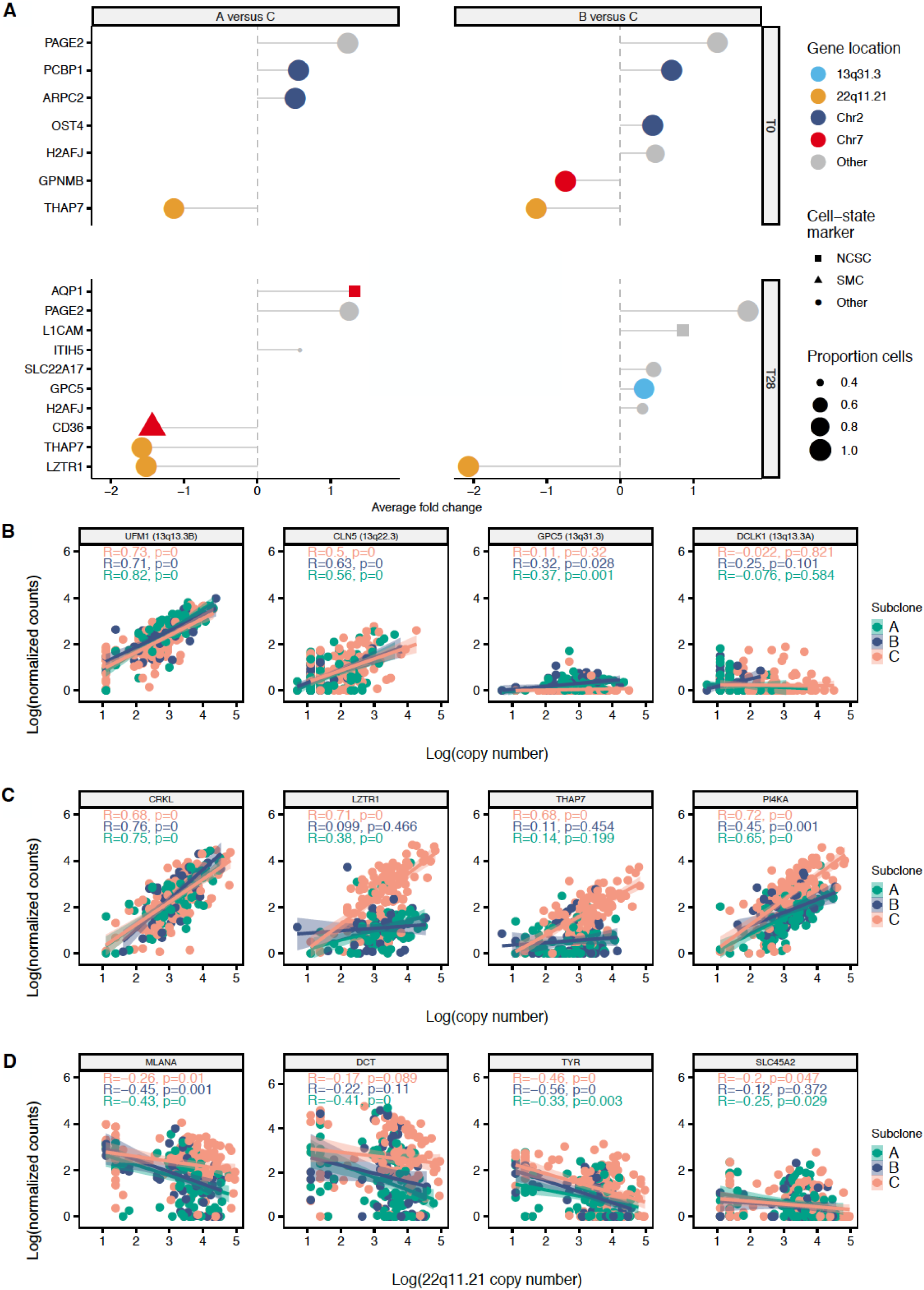
Differential gene expression analysis between subclones reveals subclone and time-point specific gene expression as well as gene dosage effects. **A** Lollipop plot depicting genes differentially expressed between genetic subclones at specific time points (FDR ≤ 0.05). No differential expression was observed between subclones A and B at either time point. The colour of the dots reflects the chromosomal location of the genes, while the shape highlights marker genes related to melanoma minimal residual disease states. The size depicts the percentage of cells expressing the gene. **B** Gene dosage plots of four genes located on the focal amplification of chr13. The copy number (log-scale with base e) of the 10 kb bin overlapping with the transcription start-site (x-axis) is plotted against log normalized gene expression counts (y-axis). **C** Same as **B** but for genes located on 22q11.21 that had observable gene expression. **D** Gene expression dosage plots for genes related to pigmentation and the copy-number of the 22q11.21 amplicon (log-scale). **B**, **C** and **D** Dots are coloured per subclone. Linear regression was performed for each subclone (shaded region indicates 95 % confidence interval). NCSC, neural crest stem cell; SMC, starved-like melanoma.

### Focal amplifications influence subclone-specific gene expression

To assess the effect of ongoing amplicon evolution on the phenotype of the subclones, we calculated the correlation between the DNA copy number of genes located on each of the focal amplifications and their normalized expression (**Fig. 4B,C** and **Supplemental Fig. S6**). For chr13 we found 5 genes with a gene-dosage effect across all the subclones (*e.g. UFM1* and *CLN5*, **Fig. 4B**), 4 genes with a subclone-specific effect (*e.g. GPC5*; **Fig. 4B**) and finally, 4 genes that did not show a dosage effect (*e.g. DCLK1;* **Fig. 4B**). All 11 genes located on the 22q11.21 amplicon showed clear gene-dosage effects (*e.g. CRKL*; **Fig. 4C**) in at least one of the subclones. Of these we found that 8 genes showed subclone-specific differences in gene expression when controlling for amplicon copy number (*e.g. LZTR1*, *THAP7, PI4KA*; **Supplemental Table S1** and **Fig. 4C**). For *LZTR1,* a tumour suppressor in many cancers although recently also suggested as a key oncogene in acral melanoma (Farshidfar et al., 2022), and *THAP7*, the disparity is elucidated by additional breakpoints in subclones A and B, resulting in a small region of lower copy number (**Supplemental Fig. S3**). The difference in *PI4KA* expression, on the other hand, did not appear to be caused by subclone-specific breakpoints. Taken together these observations suggest concomitant gene-dosage effects and epigenomic regulation modulating differential expression of genes between subclones.

### Amplicon copy number is associated with drug-tolerant cell state plasticity

Subsequently, we investigated if the amplicons were associated with phenotypic MRD states. No significant effects were observed for the majority of chr13 amplicons (**Supplemental Fig. S7A,B**). However, for the 22q11.21 amplicon we found that the expression of *MLANA*, a marker for the pigmented state, was inversely correlated with the copy number (**Fig. 4D**). In addition, genome-wide gene set enrichment analysis (GSEA) revealed a strong enrichment of gene ontology terms related to pigmentation (*e.g.* pigmentation, melanin metabolic process, pigment granule; **Supplemental Table S2**) for genes negatively correlated with the copy number of this amplicon (*e.g. DCT*, *TYR, SLC45A2*). These findings might indicate that cells with a lower 22q11.21 copy number could be more prone to occupy the differentiated pigmented cell state. In contrast, we observed a significant enrichment for high 22q11.21 copy number in the invasive state compared to both the SMC and NCSC state (**Supplemental Fig. S7D**). While there was no difference in amplicon copy number between the SMC and NCSC state, 6 genes (*e.g. KLHL22*, *TMEM191A*, *PI4KA*, *LZTR1*, *THAP7*, *TUBA3FP*) were expressed more strongly in the SMC state (**Supplemental Fig. S7C**). In summary, while none of the previously described MRD-specific markers or known melanoma-specific transcription factors are located on the 22q11.21 amplicon, our data suggests a correlation between the amplicon and the MRD states available to the subclones (Rambow et al., 2018).

### Cellular plasticity and phenotypic cell-state diversity within and between different genetic subclones on treatment

G(tag)&T-seq allows to construct, with single-cell resolution, a phylogenetic tree throughout therapy, annotated with the aforementioned drug-tolerant states. We found that while all genetic subclones were observed at both time points, their relative abundance shifted. T0 was enriched for subclones A and B (50% and 28% of cells, respectively), while subclone C was the most abundant at T28, increasing from 22% at T0 to 66% (*X*^2^-test *p* = 1.773e-13; **Fig. 5A**). Strikingly, we found the NCSC state to be enriched in subclones A and B at T28, while the SMC state was present in all genetic subclones. Both observations were validated using inferCNV and our trained SVM classifier on the Smart-seq2 data from Rambow *et al*. (Rambow et al., 2018) (**Fig. 5B**). Differential expression analysis indeed revealed higher expression of the SMC-marker *CD36* in subclone C, and increased expression of NCSC markers *AQP1* and *L1CAM* in subclones A and B respectively (**Fig. 4A**). Furthermore, GSEA revealed enrichment of genes related to epithelial-mesenchymal transition (EMT) at T28 in both subclone A and B, but not in subclone C (A, *p* = 0.0006; B, *p* = 0.0006; **Supplemental Table S3,4**). Moreover, we found that at T28, the copy number of the 22q11.21 amplicon in subclone C had increased compared to T0 (**Fig. 5C**), and to A and B at T28 (**Fig. 5D**).

**Figure 5.**
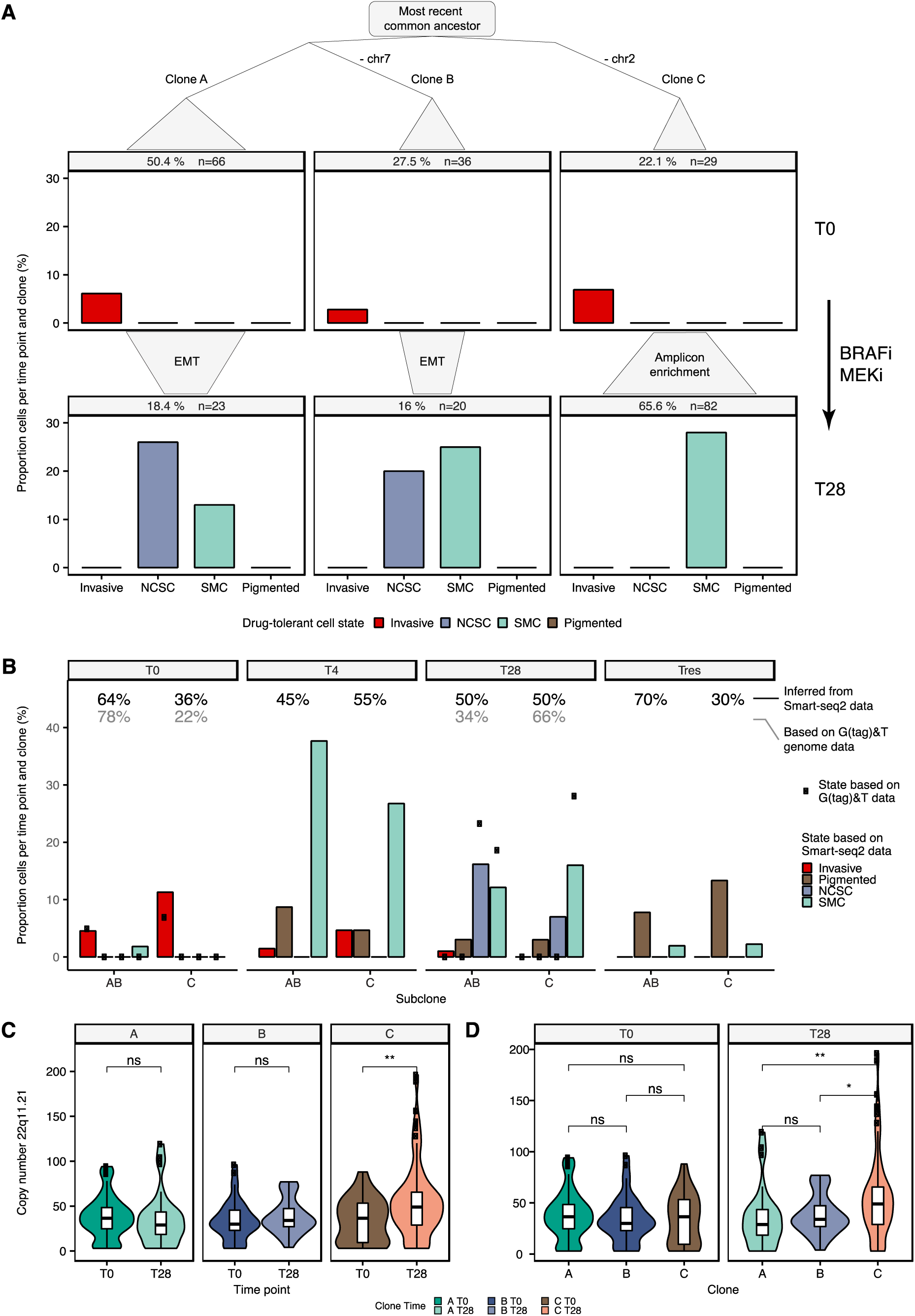
Phylogenetic tree reveals subclonal differences in treatment response and cellular plasticity. **A** Phylogenetic tree annotated with drug-tolerant cell states at T0 and T28. The shift in the proportion of cells per subclone that occurs between T0 and T28 is visualized with polygons. Gene set enrichment analysis revealed epithelial-mesenchymal transition (EMT) at T28 in subclones A and B. **B** InferCNV together with the trained SVM classifier applied to Smart-seq2 data supports the subclonal differences that were observed with G(tag)&T-seq. Subclone A and B are taken together and compared with subclone C. Percentages indicate the proportion of cells per subclone in the Gtag&T/G&T data (grey) and Smart-seq2 data (black) of Rambow *et al*. (Rambow et al., 2018). Dots represent the percentage of cells with a particular MRD state in the combined G(tag)&T-Seq data. **C** Comparison of DNA copy number distribution of the 22q11.21 amplification between timepoints (T0 *versus* T28) per subclones. **D** The same as **c**, but here subclones are compared before treatment (T0) and at MRD (T28). The centreline, top and bottom of the boxplots represent respectively the median, 25th and 75th percentile and whiskers are 1.5 × IQR. Significance levels after Wilcoxon test are as follows, ns: p > 0.05; *: p <= 0.05; **: p <= 0.01.

Taken together, we show that subclone C likely has a higher prevalence at MRD, while the NCSC state is primarily found in subclones A and B. Since drug-tolerant states appear only in a small subset of cells, direct single-cell genome-and-transcriptome methodologies are best-suited for studying this process.

## DISCUSSION

We developed Gtag&T-seq, a genome-and-transcriptome sequencing protocol of the same single cell that omits WGA by using direct genomic tagmentation. Compared to G&T-seq using picoPlex, Gtag&T-seq is characterized by improved coverage breadth and uniformity in both single-cell and pseudo-bulk genomes, which allows for more precise detection of genomic alterations. This is in line with previous reports of other research groups that used tagmentation-based library preparation without WGA (Laks et al., 2019; Rohrback et al., 2018; Vitak et al., 2017; Xi et al., 2017; Zahn et al., 2017; Zachariadis et al., 2020). Furthermore, Gtag&T-seq requires less processing time and significantly reduces the cost of G&T-seq. Moreover, it proved to be superior in the profiling of small focal amplifications (0.12-1.20 Mb), allowing breakpoints to be pinpointed to near base-pair resolution in pseudo-bulk genomes. In addition, Gtag&T allowed us to confirm the resulting gene expression dosage effects using the mRNA of the same cell.

Importantly, we highlight the need for direct multi-omics approaches to accurately dissect both genomic evolution and transcriptome plasticity in full, as opposed to inferring CNAs from single-cell transcriptomes. Besides failing to detect small focal amplifications, several whole-chromosome gains were found as smaller regions of amplification, suggesting that not all genes are affected to the same degree by genomic imbalances. Furthermore, this approach had difficulties in obtaining the correct copy number, potentially underestimating the heterogeneity present in the sample when subclones have shared breakpoints.

We applied Gtag&T-seq to a human PDX melanoma model to study the interplay of genomic and transcriptomic alterations in the context of tumour evolution and therapy resistance. Previously this model was used to identify four inducible drug-tolerant cell states –SMC, NCSC, pigmented and invasive– and revealed limited genomic heterogeneity (Rambow et al., 2018). Here, we discerned three major genomic subclones and constructed a longitudinal cell lineage tree annotated with the drug-tolerant states throughout therapy. We found subclonal differences with regard to treatment response and transcriptome plasticity: subclone C, which lost a copy of chr2, was better suited to survive the initial treatment, but had no cells with the NCSC state. In the G(tag)&T-seq data, no cells were assigned to the pigmented state while at T28 about 3% of Smart-seq2 samples from Rambow *et al*. (Rambow et al., 2018) are assigned to this state. It is unclear if the differences we observe between G(tag)&T-seq and Smart-seq2–regarding the proportion of drug-tolerant states is due to the limited number of cells analysed. In addition, we suggest a potential role for the 22q11.21 amplification in determining phenotypic differences between the subclones as well as the drug-tolerant states. One hypothesis is that this effect is mediated by *THAP7*, which is expressed higher in both subclone C and the SMC state, and is known to promote cell proliferation in lung adenocarcinoma (Chen et al., 2019). Another key gene to explain the subclonal differences could be *LZTR1* which is higher expressed in clone C cells with a high copy number of 22q11.21 and is located on a breakpoint in clone A and B. Although *LZTR1* is generally considered a tumour suppressor, a recent study in acral melanoma suggests that *LZTR1*, as well as *CRKL*, also have tumour-promoting and metastasizing capabilities (Farshidfar et al., 2022).

In the melanoma model we observed extensive heterogeneity regarding the presence, size, and dosage of focal amplicons that would be difficult to resolve using bulk sequencing and was also not always detected by picoPlex. Similar levels of amplicon heterogeneity were recently also observed in breast cancer by single-nucleus sequencing (Baslan et al., 2020). It should be further investigated whether focal amplifications, as found in the melanoma model, are drivers of subclonal differences regarding tumour evolution and therapy resistance (Francis et al., 2014), or whether they are passenger events. Nevertheless, recent studies have shown the importance of profiling driver amplicons as they proved to be predictors for survival as well as actionable targets for cancer therapy (Koboldt et al., 2012; Leucci et al., 2016; Xue et al., 2017). In this work, we show that Gtag&T is a suitable method to profile focal amplifications at near base-pair resolution in single cells.

The past years, several multi-omics techniques that interrogate the genome and transcriptome have been developed. Some of these techniques rely on separating the nucleus from the cytoplasm and are therefore limited in that nuclear mRNAs are not sequenced (Han et al., 2014; van Strijp et al., 2017; Zachariadis et al., 2020). The separation principle of transcriptogenomics (Li et al., 2015) is similar to that of G&T-seq but Li *et al*. (Li et al., 2015) only performed exome sequencing. In DR-seq (Dey et al., 2015), gDNA and mRNA are preamplified before splitting the reaction, which minimizes the risk of losing nucleic acids during the separation process. However, separating the mRNA and gDNA *prior* to amplification enables more flexibility in choosing which assay is used downstream. For example, in Gtag&T, Smart-seq2 can be replaced with the recently developed alternatives, like Smart-seq3 (Hagemann-Jensen et al., 2020), or 3’ RNA-seq technologies (Hashimshony et al., 2016; Sasagawa et al., 2018) to further reduce costs. In terms of throughput, the recently developed Sci-L3-RNA/DNA co-assay enables at least 10,000s of single nuclei to be profiled per 2-day experiment. Yin *et al*. (Yin et al., 2019) show that this combinatorial-indexing based co-assay is able to distinguish female HEK293T cells from male BJ cells based on Y chromosome presence. At the moment it is unclear how sci-L3-RNA/DNA compares to other genome-and-transcriptome technologies in terms of detected genes, and precision for detecting smaller CNAs. We do want to note that there is need for both high throughput methods, like sci-L3, as well as low-to-medium throughput when studying rare cells that can be isolated (Demeulemeester et al., 2016) or when only smaller populations of cells are available, like for example when studying genomic instability during preimplantation embryo development (Petropoulos et al., 2016; Vanneste et al., 2009).

Taken together, Gtag&T will enable researchers to study the interplay of genome and transcriptome at unprecedented detail. We suspect that our method will be broadly applicable in studying the role of somatic variation in health and disease in fields such as oncology, neurology, and embryology.

## METHODS

### Data accession codes

Human data are available from the European Genome-phenome Archive (EGA) with accession number EGAS00001007043.

### Patient-derived xenograft (PDX) model

The MEL006 cutaneous melanoma PDX model is part of the Trace collection (https://gbiomed.kuleuven.be/english/research/50488876/54502087/Trace/PDX-repository) and was established using lesions derived from a patient undergoing surgery as part of standard treatment at UZ Leuven. Written informed consent was obtained and all procedures were approved by the UZ Leuven/KU Leuven Medical Ethical Committee (S63799, S57760, S58277) and performed in accordance with the principles of the Declaration of Helsinki and with GDPR regulations. The experiments were approved by the KU Leuven animal ethical committee under ECDs P164-2019 and performed in accordance with the internal, national, and European guidelines of animal care and use. Single-cell suspensions were implanted subcutaneously in the interscapular fat pad of female NMRI nude BomTac:NMRI-Foxn1nu, 4-wk-old females (Taconic Biosciences). Mice were maintained in a pathogen–free facility under standard housing conditions with continuous access to food and water. The health and welfare of the animals was supervised by a designated veterinarian. The KU Leuven animal facilities comply with all appropriate standards (cages, space per animal, temperature [22°C], light, humidity, food, and water), and all cages are enriched with materials that allow the animals to exert their natural behaviour. Mice used in the study were maintained on a diurnal 12-h light/dark cycle. MEL006 was derived from a female, drug-naïve melanoma patient. When the tumour reached 1000 mm^3^ the mice were randomly assigned to the different experimental groups. Mice were treated daily by oral gavage with a capped dose of 600 μg dabrafenib –6 μg trametinib (DT), in 250 μl total volume.

### Sample preparation for single-cell sorting

HCC38 breast cancer cells and HCC38-BL lymphoblastoid cells were cultured in Dulbecco’s Modified Eagle Medium (DMEM/F12) containing 10% fetal bovine serum (FBS) at 37°C in a 5% CO2 incubator (Voet et al., 2013). Trypsinized HCC38 cells and HCC38-BL cells were washed in fresh culture medium and then resuspended in FACS sorting buffer (DMEM/F12 supplemented with 5% FBS, 1mM EDTA, and 1.5 μM DAPI). The BD FACS Melody sorter device was used for sorting single cells into 96-well plates (FrameStar®, 4TI-0960/C) containing 2.5 μl RLT plus buffer (Qiagen). Plates were then spun down at 1,000g for 1 minute at 4°C, and finally stored at −80 °C. MEL006 cells were retrieved as described before (Rambow et al., 2018), resuspended in serum-free DMEM/F12 medium and sorted using the BD FACS Aria III into 96-well plates containing 2.5 μl RLT plus buffer.

### Gtag&T

Genomic DNA and mRNA separation and DNA precipitation was performed as per the G&T protocol of Macaulay *et al*. (Macaulay et al., 2015, 2016) on an Hamilton^®^ liquid handling robot. First, the plate containing the lysed cells is supplemented with RNA spike-ins (1 μL of a 1:1,600,000 dilution of ERCC spike-in mixture A (Life Technologies)), which is followed by the addition of oligo-dT conjugated to streptavidin beads. After incubation, the poly-adenylated mRNAs are collected to the side of the well using a magnet, while the DNA, present in the supernatant, is transferred to a new recipient DNA plate. The wash solution used to rinse the mRNA-bead complexes is also added to the DNA plate. After mixing the solution containing the DNA with Ampure XP beads, the DNA is precipitated using a magnet. For Gtag, the precipitated genomic DNA was resuspended in 4.5 μl resuspension buffer (0.5X NEB4, 0.37% Igepal CA-630, 0.37% Tween-20). Next, tagmentation is performed in a total volume of 10 μl by adding 5.5 μl tagmentation master mix (5 μl Tagment DNA buffer, 0.1 μl Tagment DNA Enzyme, 0.4 μl nuclease free water) for 10 minutes at 55°C. The reaction was inactivated by adding 1 μl of 0.44 % SDS to the sample and incubating 5 minutes at 55°C. Then we added 13 μl of Q5 Ultra II (NEB, 2x mastermix), 1 μl S5 primer, and 1 μl S7 primer to the sample. PCR amplification was performed with the following cycling program: 72°C for 3 min; 98°C for 30 sec; 16 cycles of 98°C for 10 sec, 60°C for 30 sec and 72°C for 30 sec; 72°C for 5 min; and held at 10°C. In between reaction steps, 96-well plates were placed in an Eppendorf thermomixer at room temperature to mix (1,000 rpm) for one minute, and briefly centrifuged using a tabletop centrifuge. Finally, the PCR products were pooled per plate and purified using AMPure XP beads (1x ratio).

### Sequencing library preparation

DNA and cDNA quality of picoPlex and Smart-seq2 amplification reactions, respectively, was confirmed using the 2100 Bioanalyzer (high sensitivity chip, Agilent). Next, DNA and cDNA concentrations were determined using a Quantifluor^®^ Assay (Promega^®^). Samples were diluted to 200 pg/μL or 100 pg/μL for Nextera XT (Illumina) library preparation in respectively one-fourth or one-tenth of the volume recommended by the manufacturer using manual or automated liquid handling. After library preparation, samples were pooled and purified using AMPure XP beads (0.6x ratio). Quality check of the DNA (picoPlex and Gtag) and cDNA library pools was performed using the 2100 Bioanalyzer (high sensitivity chip, Agilent) in combination with Qubit™ HS (High sensitivity) DNA Assay Kit (Invitrogen™) before diluting the pools to a concentration of 4nM. We used the KAPA Library Quantification Kit for Illumina^®^ platforms (Roche, KK4854) on the LightCycler 480 and diluted the pools to 2nM (HCC38 samples) or 1.5 nM (MEL006 samples) before sequencing 51-bp single-end reads on HiSeq2500 or HiSeq4000 respectively.

### Processing of genome data and DNA copy number analysis

Single-end sequencing reads obtained were aligned to the GRCh37 human reference genome using BWA-MEM (Li, 2013). Samtools was used to sort, index, and sample the mapped BAM files down to 400,000 reads. Our mapping statistics were obtained through samtools and Picard (http://broadinstitute.github.io/picard/). PCR duplicates were removed with Picard. To create pseudo-bulk genomes, *samtools merge* was used to combine BAM files. DNA copy number analysis was performed as discussed in Macaulay *et al*. (Macaulay et al., 2015, 2016). Segmentation of the corrected logR values was done using piecewise constant fitting, with the penalty parameter (γ) set to 10 for the 500 kb bin genomes and γ = 35 for the 10 kb bin genomes. The average ploidy was chosen based on logR values of a large region with few to no copy number aberrations:

| Sample | Avg ploidy | Ref region |
| --- | --- | --- |
| HCC38 | 3 | chr12 |
| HCC38-BL | 2 | all chr |
| MEL006 | 3 | chr3 |

### scDNA-seq quality filtering

For quality filtering of MEL006 genomes, we only processed single cells with at least 400,000 reads before deduplication. HCC38 genomes were only processed if they had at least 100,000 raw reads before deduplication. We calculated the Median Absolute Pairwise Difference (MAPD) score for all samples by first measuring the absolute difference between two consecutive logR values, %GC-corrected and normalized, across the genome. Next, the median across all absolute differences is computed. For MAPD cut offs, genomes only passed if their MAPD score was less than the 75^th^ percentile + 1.5 times the interquartile range (HCC38, MAPD_cut off_ = 0.64; MEL006, MAPD_cut off_ = 0.69). An overview of the QC pass/fail samples can be found in **Supplemental Data S1** and **S2**.

### Coverage uniformity calculations

Lorenz curves were computed by taking the cumulative fraction of the covered genome against the cumulative fraction of the mapped bases. From the BAM files, duplicates were first removed, and all genomes were down-sampled to 230,000 unique reads (with a quality of at least 20). Gini coefficients were calculated in R using the ineq package (https://cran.r-project.org/web/packages/ineq/).

### Heatmaps and clustering

The Canberra distance was used as a dissimilarity metric between the DNA copy number profiles of individual single cells. Hierarchical clustering was performed using Ward’s D2 criterion to minimize the total variance within each cluster. Both the gap statistic and the silhouette score indicated the two as the optimal number of clusters, resulting in the detection of subclones A|B and C. However, the generation of pseudo bulks for these clusters, as well as manual inspection revealed remaining heterogeneity regarding chr7 in cluster A|B. Therefore, we extended the number of clusters to five, wherefore no remaining heterogeneity was observed in the pseudo-bulk genomes. Regions of focal amplifications were determined on the pseudo-bulk Gtag genomes on 10 kb bins segmented by piecewise constant fitting (γ = 35). For each region, 20 additional genomic bins were taken on each flank for visualization. All heatmaps were constructed using the ComplexHeatmap package (version 2.2.0).

### Processing of single-cell RNA seq data

After trimming of the adaptor sequences with cutadapt (version 1.13), sequencing reads were aligned to the GRCh37 reference genome including ERCC sequences using STAR with default parameters (version 2.5.2b). HTseq (version 0.6.0) with the GENCODE H19 transcript annotations were used to generate the count matrix.

### Analysis of single-cell RNA-seq: HCC38 and HCC38-BL

Quality control was performed using the scater R package (version 1.10.1): cells with less than 100,000 counts, expression of less than 2000 unique genes, more than 30% counts assigned to mitochondrial sequences or 8% counts belonging to ERCC sequences were removed for downstream analysis. Genes with less than 32 counts across the complete dataset were excluded from downstream analysis. All data analysis was conducted in R v3.5.0 (CRAN), while plots were created with the ggplot2 (version 3.0.0) R package.

### Analysis of single-cell RNA-seq: MEL006

Quality control was performed using the scater R package (version 1.10.1): cells with less than 100,000 counts, expression of less than 1000 unique genes, more than 25% counts assigned to mitochondrial sequences or 15% counts belonging to ERCC sequences were removed for downstream analysis (McCarthy et al., 2017). Genes with less than 5 counts across the complete dataset were excluded from downstream analysis. Expression value scaling and normalization, cell-cycle regression, batch correction, PCA and UMAP dimensionality reductions and clustering were performed using the Seurat R package (version 3.0.2) (Butler et al., 2018). Marker gene discovery was performed using the FindAllMarkers function of the Seurat package using the Wilcoxon Ranked Sum test. A clustering resolution of 0.4 was selected since it revealed clusters with biological meaningful significance as well as displaying cluster stability when using the clustree R package (version 0.4.0). pySCENIC (commit 0.9.9+2.gcaded79) (Aibar et al., 2017) with default parameters was used to perform gene set regulatory network analysis. The R package presto (version 1.0.0) was used to perform a fast Wilcoxon rank sum test where the AUC value served as input for gene set enrichment analysis with FGSEA (version 1.8.0.). Gene sets were accessed with msigdbr (version 6.1.1). Gene set enrichment analysis was also performed with FGSEA using the correlation scores of all genes passing quality control and the copy number of the 22q11.21 amplicon. Gene dosage plots were created for all genes located on focal amplifications when at least 10 single cells expressed the gene, the unsegmented copy number was taken from the bin overlapping with the transcription start site for each gene. The copy number of the bin closest to the middle of focal amplification was used as the overall copy number of the amplicon. MRD states were assigned as described in Rambow *et al*. (Rambow et al., 2018). All data analysis was conducted in Python v3.6 (Python software foundation) or R v3.5.0 (CRAN). Plots were created with the ggplot2 (version 3.0.0) and ggpubr (version 0.2.1) R packages.

### Benchmarking of inferCNV with G(tag)&T-seq data and classification of the Rambow *et al*. (Rambow et al., 2018) Smart-seq2 data

Raw single-cell RNA-seq reads from 96 normal human melanocytes (ethics approval S63257) (data available on request), as well as the Rambow *et al*. (Rambow et al., 2018) (GEO: GSE116237) data were aligned to the GRCh37 reference genome including ERCC sequences using STAR with default parameters (version 2.5.2b). After creating a count matrix with HTseq (version 0.6.0) and the GENCODE H19 transcript annotations, the data was merged with the G(tag)&T transcriptome counts. The scater R package was used to discard low-quality cells, namely cells with less than 100,000 counts, less than 1000 unique genes expressed, more than 25% counts assigned to mitochondrial sequences or more than 15% counts belonging to ERCC sequences. InferCNV v1.2.1 was subsequently used to infer copy number estimates from the scRNA-seq data using standard parameters for Smart-seq2 and the 6-state Hidden-Markov Model while using the transcriptome data of the normal melanocytes as reference. The mitochondrial and sex chromosomes were excluded from the analysis. The G(tag)&T data was used to benchmark the copy number calls obtained with inferCNV, where the DNA-seq data was considered the gold truth. Segments were then classified as (i) true positive if both DNA and RNA copy number calls indicated a gain or if both indicated a loss; (ii) true negative if both indicated a neutral copy number state; (iii) false negative if the DNA data indicated a copy number aberration and the RNA did not; (iv) false positive if the RNA indicated a copy number aberration and the DNA did not. The copy number calls, and modified expression intensities obtained with inferCNV for the G(tag)&T transcriptome data were both used to train 5 machine learning algorithms (Linear Discriminant Analysis, Classification and Regression Trees, k-Nearest Neighbours, radial function Support Vector Machine (SVM) and Random Forest) to classify a sample to the correct genomic subclone. Briefly, the caret R package (Version 6.0-80) was used to split the data 80%-20% after removing highly correlated features (cor > 0.8) and train the classifiers using 10-fold cross-validation. Accuracy was selected as the scoring metric to assess the performance. The best accuracy on the test data was observed for the radial function SVM trained on the modified expression intensities when combining subclones A and B. The model was then used to assign the Rambow et al. (2018) to either subclone AB or C. Data analysis was conducted in R v3.5.0 (CRAN) while plots were created with the ggplot2 (version 3.0.0) and ggpubr (version 0.2.1) R packages.

## ACKNOWLEDGEMENTS

The authors would like to thank the Genomics Core Leuven for sequencing, FACS Core Leuven for sorting, S. Geurs and I. Smeers for reading the manuscript. We would like to acknowledge our colleagues from the Sanger Single Cell Genomics Core Facility and in particular: Dr. Stephan Lorenz and Dr. Iraad Bronner. S.V. and K.T. are supported by a FWO (Research Foundation – Flanders) PhD fellowship for Strategic Basic Research (1S93318N and 1126016N). K.T. is supported by a Kom op tegen Kanker starters grant (ZKC8518). J.D. and A.S. are supported by a postdoctoral fellowship of the FWO (12J6916N and 12W7318N). T.V. is supported by KU Leuven (SymBioSys - C14/18/092), Foundation Against Cancer (2015-143) and FWO (I001818N). O.M-B. is supported by 12T1217N project by FWO at the program under the Marie Skłodowska-Curie grant agreement no. 665501. N.V.R. is supported by the FWO Strategic Basic research grant 1S79619N. F.R. received postdoctoral research fellowships from the Omics/Marie Curie at VIB. J.-C.M. is supported by grants from the Melanoma Research Alliance Established Investigator Award (#623591), FWO (G097918N), KULeuven (C1, C16/19/006). E.L. is supported by grants from Melanoma Research Alliance young investigator award 2018 (https://doi.org/10.48050/pc.gr.80542), from KULeuven (C1, C16/19/006). Trace Platform is supported by Stichting Tegen Kanker grant 2016-054.

## Author contributions

K.T., S.V. and T.V. designed the experiments and performed data analysis. K.T., S.V., J.D. and T.V discussed the data and wrote the manuscript. K.T. developed Gtag&T with help of D.B.. S.V. performed the bioinformatic analysis with help of K.T. and M.V.D.H, A.S. and F.R.. A.R, O.M.B., N.V.R., E.L. and J.M. provided the normal melanocytes and human melanoma PDX model. All authors read and approved the final manuscript.

