## Supplemental_Figures_Data_Tables for "Single-cell Genome-and-Transcriptome sequencing without upfront whole-genome amplification reveals cell state plasticity of melanoma subclones": Supplemental_Figures.pdf

Supplemental Fig. S1. RNA and DNA sequencing statistics of G&T-seq versus Gtag&T-seq for the HCC38 BL cell line and HCC38 cell line.

Supplemental Fig. S2. MEL006 DNA and RNA sequencing statistics.

Supplemental Fig. S3. Subclonal 20 kb alterations are not detected in single-cell picoPlex genomes.

Supplemental Fig. S4. The focal amplifications of chr13 correlate with amplification of 22q11.21.

Supplemental Fig. S5. Focal amplifications display genetic heterogeneity in presence, size and copy number.

Supplemental Fig. S6. Gene dosage plots reveal subclone-specific gene expression.

Supplemental Fig. S7. Cells from the invasive state show higher amplicon copy number compared to other drug tolerant cell states.

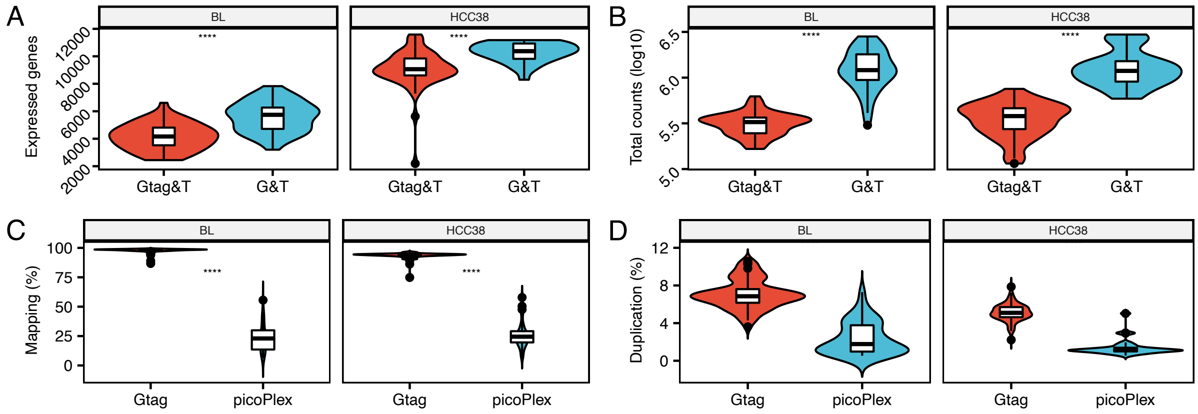


**Supplemental Figure S1. RNA and DNA sequencing statistics of G&T-seq *versus* Gtag&T-seq for the HCC38 BL and HCC38 cell line.** Comparison of **A** the number of expressed genes, **B** total genomic counts, **C** the amount of genomic mapped reads, and **D** the percentage of genomic PCR duplicates.

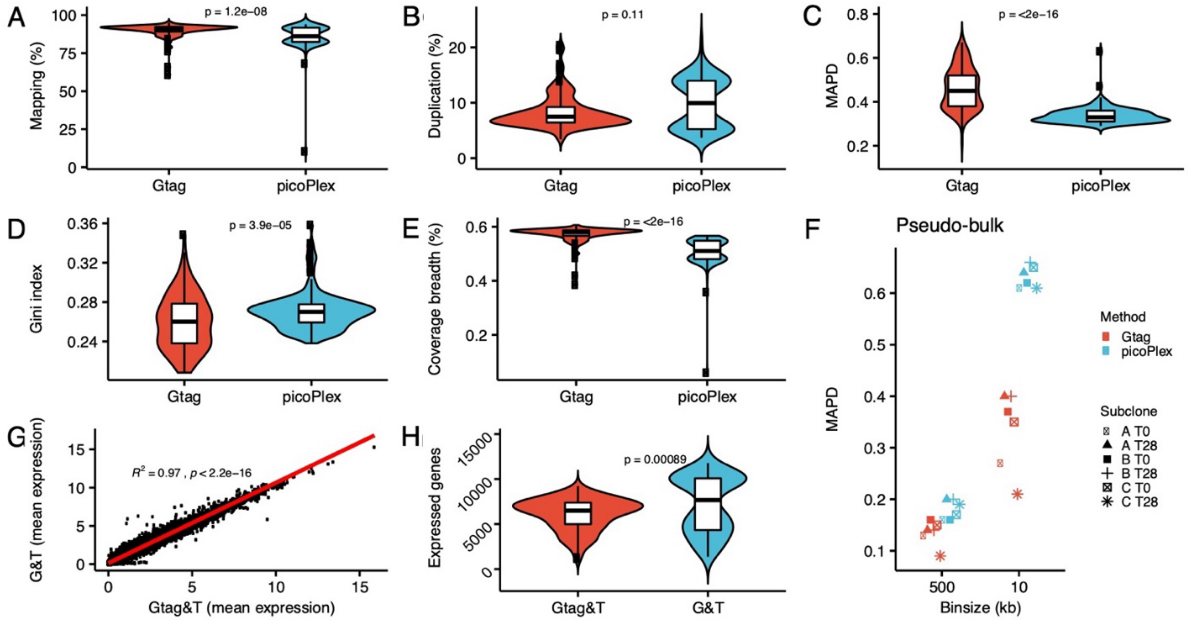


**Supplemental Figure S2. MEL006 DNA and RNA sequencing statistics.** Quality control of DNA samples in **A**-**E**. Comparison of the number of mapped reads (**A**), the percentage of PCR duplicates (**B**). Although MAPD values were overall lower for picoPlex genomes (**C**), Gtag remained superior in coverage uniformity (**D**) and breadth of coverage (**E**). The maximally attainable coverage at this depth is 0.667 %. **F** Gtag pseudo-bulk genomes obtained better MAPD values. **G** Spearman correlation of the mean expression per gene, and **H** the number of expressed genes.

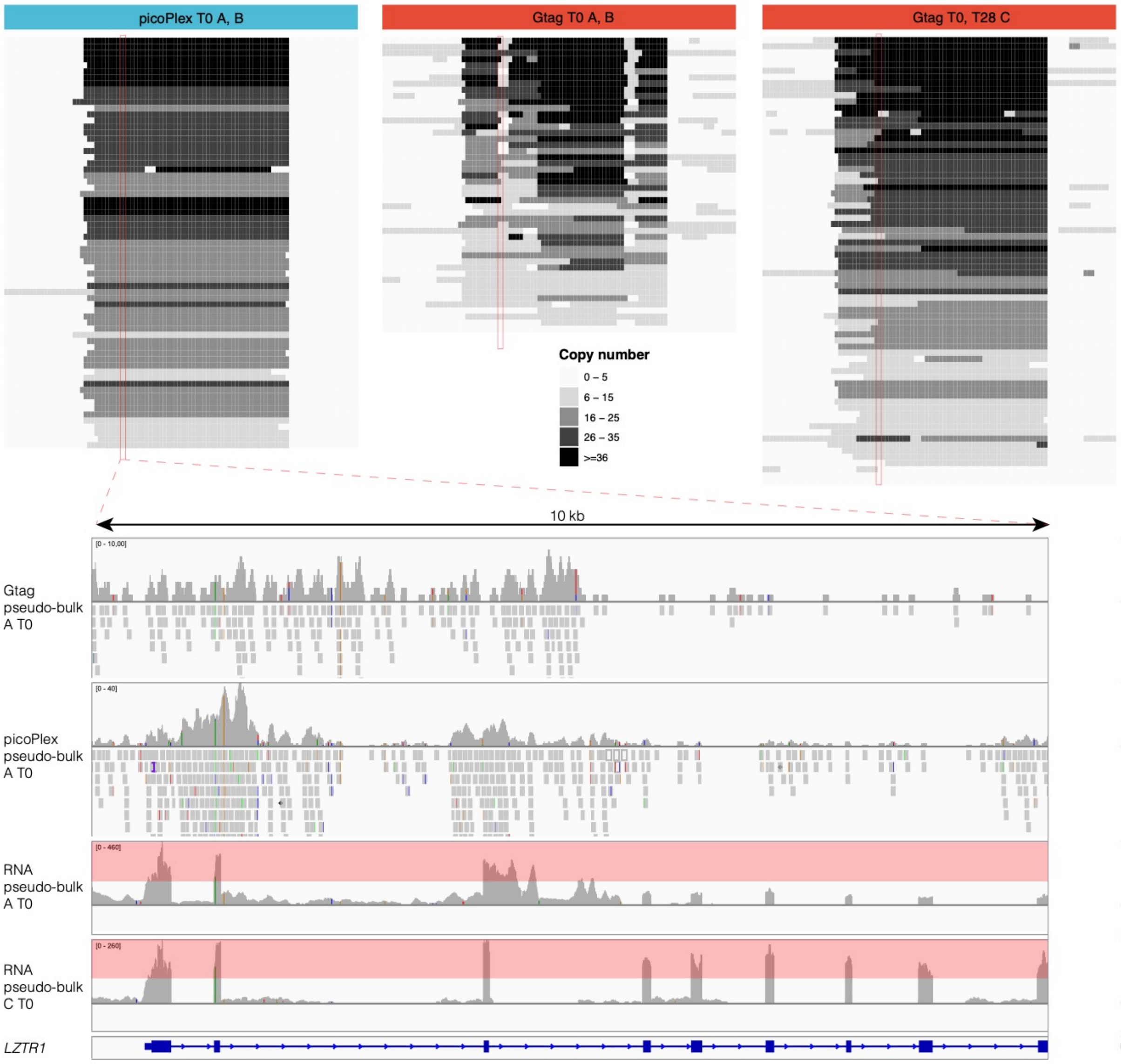


**Supplemental Figure S3. Subclonal 20 kb alterations are not detected in single-cell picoPlex genomes.** Upper panels show the 22q11.21 amplicon for single-cell genomes, analyzed with 10 kb bins, from picoPlex and Gtag (subclone A and B at T0) and Gtag (subclone C at T0 and T28). Red rectangles (10 kb window) indicate the location of the breakpoint that is not detected in picoPlex genomes. The lower panels represent an IGV view of the same window for DNA pseudo-bulk genomes and RNA pseudo-bulks. Exons downstream from the breakpoint show less expression hereby validating the alteration.

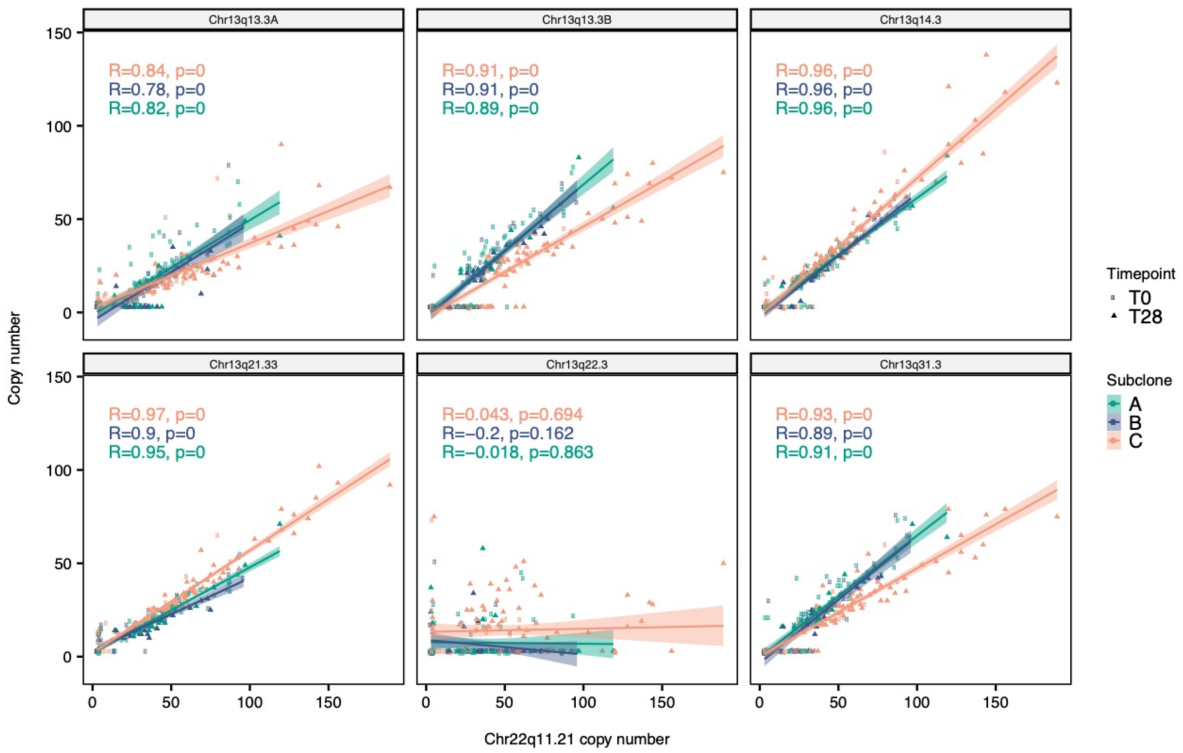


**Supplemental Figure S4. The focal amplifications of chr13 correlate with amplification of 22q11.21.** Correlation of the copy number of 22q11.21 (x-axis) with the copy number of six amplicons from chr13 (y-axis). The specific amplicon of chr13 is indicated in the header boxes. The copy number of the 10 kb bin closest to the middle of the amplicon was used as the overall copy number for the focal amplifications.

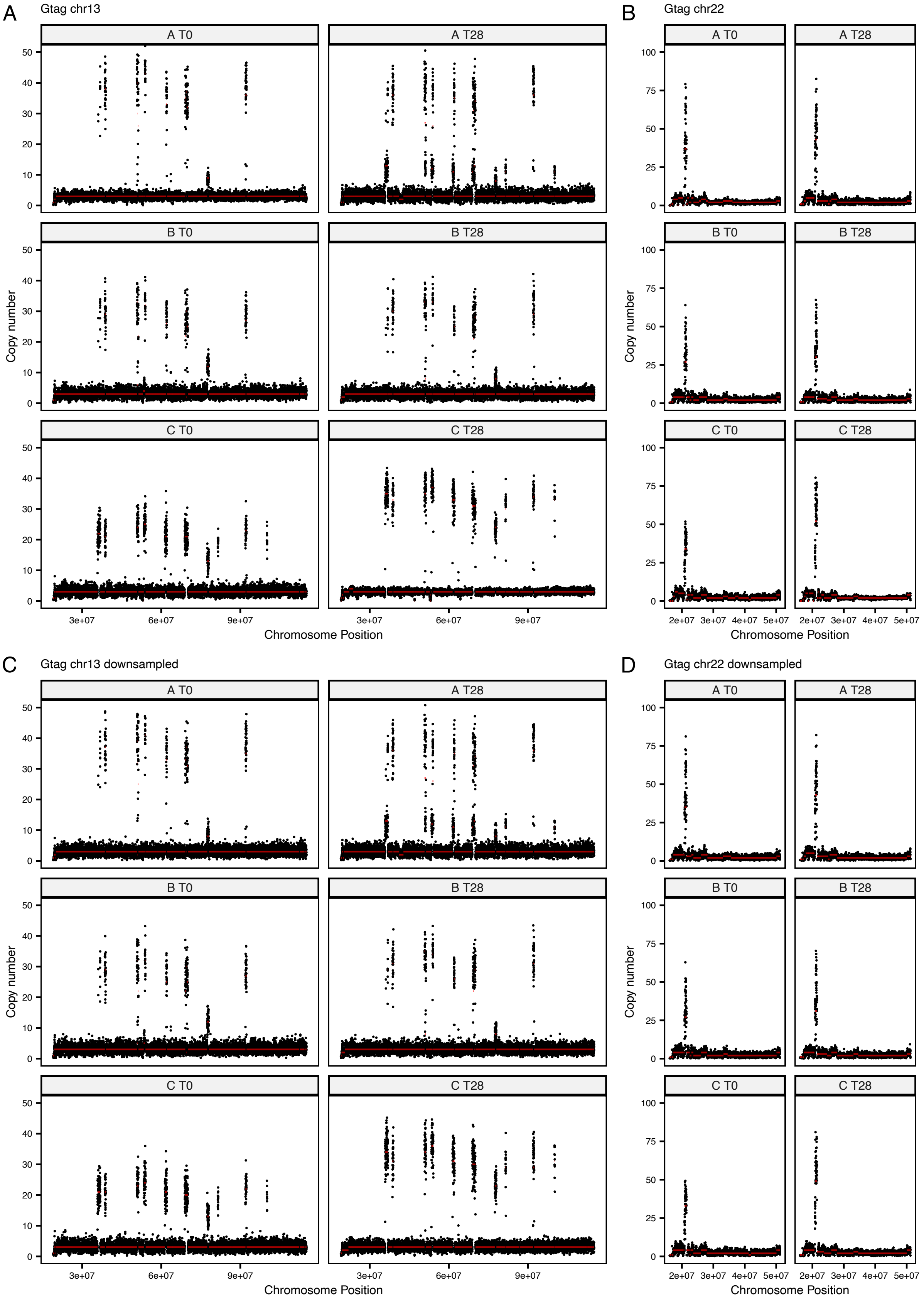
A
Gtag chr13
B
Gtag chr22
C
Gtag chr13 downsampled
D
Gtag chr22 downsampled
A T0
A T28
B T0
B T28
C T0
C T28
Copy number
Chromosome Position
3e+07
6e+07
9e+07
2e+07
3e+07
4e+07
5e+07

**Supplemental Figure S5. Focal amplifications display genetic heterogeneity in presence, size and copy number.** **A** DNA copy number profile of chr13 from Gtag pseudo-bulk genomes at T0 and T28. **B** Same as **A** for chr22. **C** and **D** show the same as respectively **A** and **B** but here all pseudo-bulk genomes are down-sampled to the same depth in order to correct for differences caused by sequencing depth.

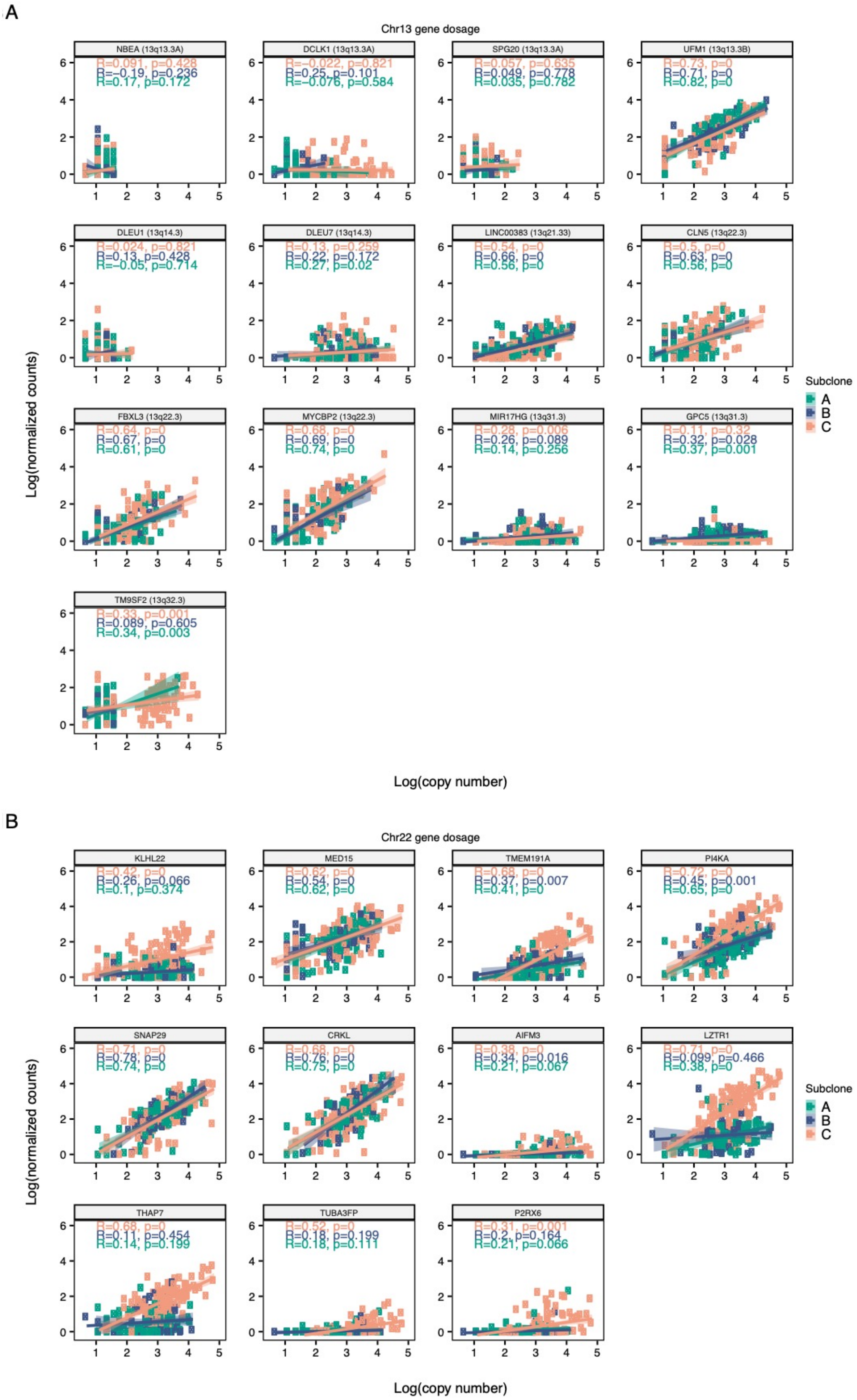


**Supplemental Figure S6. Gene dosage plots reveal subclone-specific gene expression. A** Gene dosage plots for all expressed genes located on the focal amplification of chr13. **B** Same as **A** but now for genes located on 22q11.21 that had observable gene expression. The copy number (log-scale) of the 10 kb bin overlapping with the transcription start-site (x-axis) is plotted against log normalized counts (y-axis). Dots are coloured per subclone. Linear regression was performed for each subclone (shaded region indicates 95 % confidence interval).

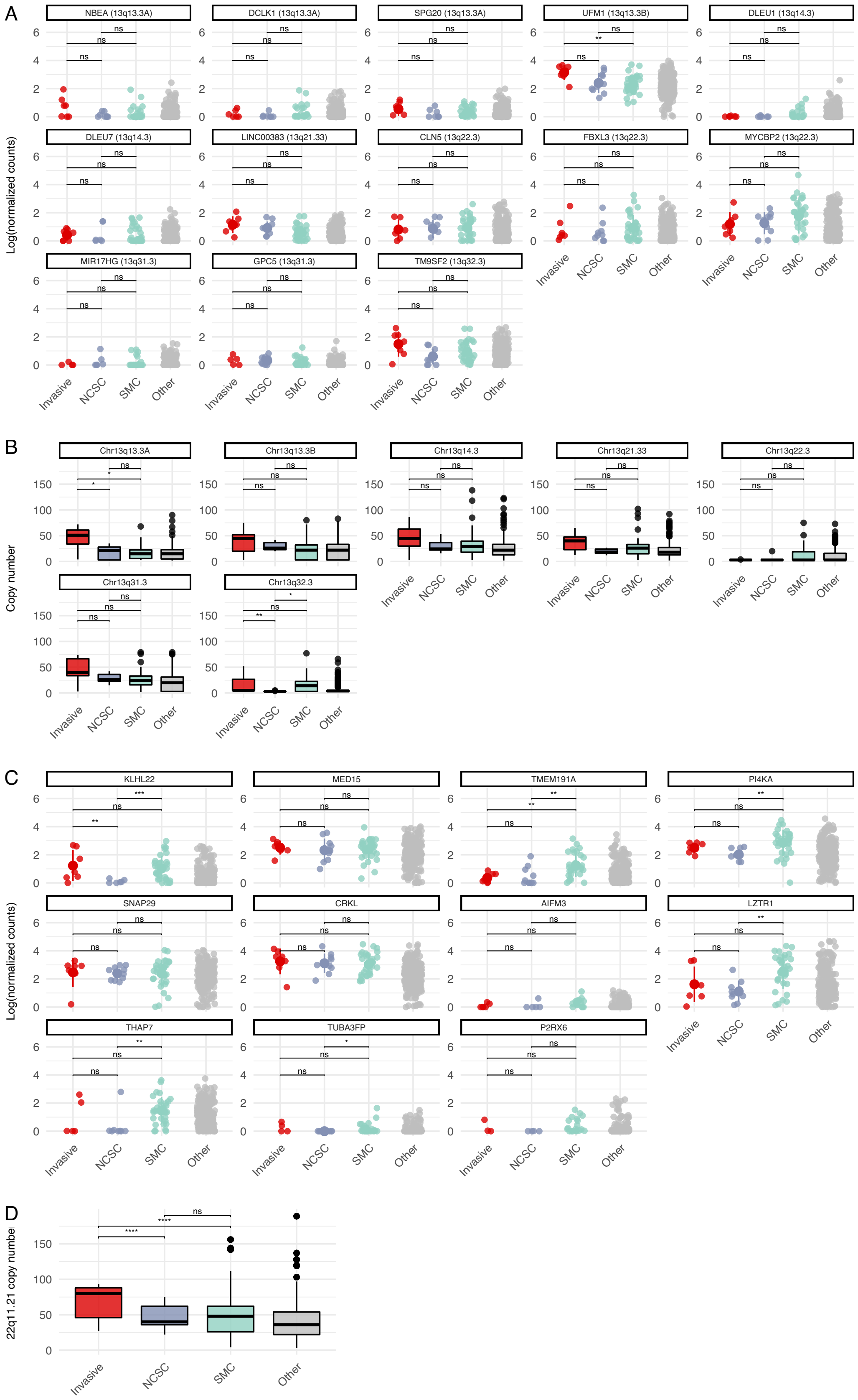


**Supplemental Figure S7. Cells from the invasive state show higher amplicon copy number compared to other drug tolerant cell states. A** One-dimensional scatter plot representing the distribution of log-normalized expression counts for the genes with

observable expression located on the focal amplifications of chr13 across the different MRD states. **B** Copy-number distribution for the focal amplifications on chr13 across the different MRD states. **C** Same as **A** but for genes located on 22q11.21 that had observable gene expression. **D** Same as **B** but for the 22q11.21 focal amplification.
The centerline, top and bottom of the boxplots represent respectively the median, 25th and 75th percentile and whiskers are 1.5 × IQR. Beginning and end of the line in b and d represent the 25th and 75th percentile with the dot indicating the median. Significance levels after Wilcoxon test are as follows, ns: $p > 0.05$; *: $p <= 0.05$; **: $p <= 0.01$. The other category contains cells not belonging to one of the MRD states.
